# SIGMA: SIgnaling via Golgi Morphology Analysis for pathway-state classification

**DOI:** 10.64898/2026.10.01.756010

**Authors:** Mengchen Fan, Xu Zhang, Baocheng Geng, Pulin Che

## Abstract

Organelle morphology is increasingly recognized as a computationally accessible readout of molecular state, yet whether subcellular architecture can identify which signaling pathway is active, not merely whether a cell is perturbed, remains untested. Here we introduce SIGMA (SIgnaling via Golgi Morphology Analysis), a framework that reframes pathway identification as a morphological classification problem centered on the Golgi apparatus. SIGMA integrates imaging flow cytometry for population-scale Golgi profiling, teacher-student knowledge distillation to transfer fine morphological cues from limited high-resolution images to an efficient lower-resolution classifier, and mechanistic generalization testing with chemically distinct activators. Applied to MRC-5 fibroblasts across six conditions spanning four signaling cascades plus controls, SIGMA achieved 90.02% cell-level accuracy and correctly assigned five of seven structurally unrelated external compounds to their expected pathway class. These results establish Golgi morphology as a multiplexed, machine-readable fingerprint of signaling state.

## Introduction

Organelle morphology and cellular function are intrinsically linked, with structural features not merely reflecting but actively contributing to functional capacity, a relationship most explicitly formalized for mitochondria as’morphofunction’ ^1^ and recognized more broadly across organelles, including ER shape determining ER function and organelle positioning controlling cellular processes ^2, 3^. This principle has been compellingly demonstrated for mitochondria, where network architecture directly influences ATP production efficiency: elongated networks correlate with high-energy states, whereas fragmented morphologies accompany mitophagy and reduced bioenergetic capacity ^4, 5^. Recent computational advances have transformed this qualitative understanding into quantitative prediction. Deep learning analysis of mitochondrial morphology now enables accurate prediction of metabolic functions including membrane potential, respiration rate, and ATP generation, achieving strong correlations between predicted and measured values ^6^. Similarly, whole-cell morphological profiles can predict signaling-pathway perturbations and drug-resistance mechanisms ^7, 8^. These findings establish that organelle and cellular architecture contain computationally extractable information about molecular functional states, raising a fundamental question: does this principle extend to organelles that integrate diverse signaling inputs, and could such relationships be exploited for practical applications including drug efficacy prediction?

The Golgi apparatus is well suited for investigating morphology-signaling relationships because it functions as both a structural organelle and a signaling integration platform ^3, 9, 10^. Unlike organelles with relatively circumscribed functions, the Golgi receives regulatory inputs from cell cycle machinery, stress response pathways, secretory demand signals, and polarity cues, integrating these diverse inputs into coherent structural states ^3, 10, 11, 12, 13, 14^. These morphological responses exhibit pathway specificity rather than generic stress reactivity: PKA activation drives ribbon compaction associated with cAMP signaling ^15^; calcium store depletion triggers PKCα-mediated GRASP55 phosphorylation and Golgi fragmentation ^16^; and PKC activation produces fragmentation through GRASP55 phosphorylation by a partially overlapping mechanism ^16, 17^. The specificity of these responses suggests that Golgi architecture might encode information about which signaling pathways are active, not merely whether the cell is perturbed. If this hypothesis proves correct, Golgi morphological analysis could provide a readout of pathway activation status with potential utility for characterizing drug mechanisms of action and predicting therapeutic efficacy.

Despite this biological foundation, methodological challenges have constrained previous investigations of Golgi morphology-signaling relationships. Conventional microscopy-based morphological analysis remains labor-intensive and suffers from limited statistical power, with typical studies analyzing dozens to low hundreds of cells ^18^. Threshold-based segmentation methods are sensitive to image quality and intensity variation, often requiring manual tuning that limits reproducibility across large datasets and motivating recent shifts toward deep-learning-based organelle segmentation ^19, 20, 21^. Most existing computational approaches to organelle image analysis have focused primarily on cell-level outcomes such as gene-knockdown phenotyping or cell-cycle stage, rather than attempting to link morphology with specific pathway activation states ^22, 23, 24, 25^. This leaves a clear methodological gap: developing approaches that quantitatively connect Golgi morphology with signaling pathway activation could advance understanding of organelle-signaling relationships while establishing practical tools for pathway characterization applicable to drug discovery and therapeutic development.

Recent advances in machine learning have changed biomedical and subcellular imaging analysis ^26, 27^. Deep learning segmentation and feature extraction approaches now allow automated, reproducible quantification of cellular and organelle morphology across large image datasets, overcoming limitations of manual or rule-based analysis ^21, 28^. Knowledge-distillation methods provide a complementary strategy for transferring information from high-capacity teacher models into efficient student networks, and recent biomedical imaging work has extended this idea through stage-wise distillation designed to preserve fine image detail ^29, 30, 31^. Beyond structural quantification, emerging studies have demonstrated that machine learning models can extract biologically meaningful information from morphology to predict functional or molecular states ^6, 7^. These capabilities highlight the potential of morphology as an information-rich readout rather than a purely descriptive phenotype. However, despite these advances, most applications remain focused on either structural segmentation or prediction of single functional outputs and rarely attempt to infer specific signaling pathway activation states from subcellular morphology. Existing approaches also typically demonstrate generalization at the level of image similarity or compound identity, without rigorous testing showing that learned representations reflect shared signaling mechanisms rather than compound-specific effects. These limitations underscore a gap of pathway-level, mechanistically validated inference frameworks capable of linking subcellular morphology to distinct signaling states.

To address these challenges, we developed SIGMA (SIgnaling via Golgi Morphology Analysis), a computational framework for Golgi image analysis and signaling pathway classification. Whereas recent computational approaches have linked organelle morphology to continuous functional outputs such as mitochondrial metabolic activity ^6^ or kinase pathway activity ^7^, SIGMA addresses the distinct problem of classifying discrete signaling pathway states from Golgi morphology. Conceptually, SIGMA treats Golgi morphology as a pathway fingerprint, a structural readout that identifies which of several signaling pathways is active rather than merely whether a cell is perturbed, extending this morphological-fingerprinting concept to a signaling-integrating organelle. The framework operates through three integrated modules designed to progress systematically from foundational biological validation through high-throughput data acquisition to predictive classification and generalization testing **(Figure 2)**. Module 1 combines imaging flow cytometry with rigorous quality control to generate morphological profiles from 33,061 raw single-cell images. Module 2 transfers fine Golgi-morphology information from limited high-resolution crops into a lower-resolution ImageStream student through teacher-guided distillation. Module 3 validates generalization through mechanistic testing with chemically distinct compounds that engage identical pathways, thereby distinguishing pathway-associated features from compound-specific artifacts. This modular architecture is designed to keep classification tied to biologically meaningful relationships between Golgi structure and signaling function.

We applied SIGMA to classify MRC-5 human lung fibroblasts across four signaling pathway activation states (PKA, JNK/p38 MAPK, PKC, and ER stress/Ca²⁺), a Wnt-based morphological negative control (CHIR99021), and vehicle, using well-characterized pharmacological perturbations as a proof-of-concept demonstration. Our results indicate that Golgi morphological patterns contain sufficient information to discriminate among these conditions with 90.02% cell-level test accuracy, that morphological feature importance varies in biologically plausible patterns across pathways, and that compound-level testing is perfect on the internal test set but partial on the external chemically distinct test panel, revealing compound-specific failure modes that define current boundaries of morphology-based pathway inference. These findings demonstrate that Golgi architecture encodes signaling-state information amenable to computational extraction, extending the principle that organelle morphology predicts molecular state to a central trafficking hub. Beyond methodological advances, this work demonstrates the potential of integrating high-throughput imaging with machine learning for subcellular structure analysis, providing proof-of-concept evidence that organelle morphological signatures can inform pathway activation status for pathway classes that produce distinct Golgi phenotypes. Such capabilities may complement conventional biochemical approaches in drug discovery and development, particularly for rapid morphology-based assessment of pathway engagement where training-data coverage is well matched to the candidate compound’s mechanism of action.

## Results

### Signaling pathway activation produces distinct, quantifiable Golgi morphologies

Classifying signaling pathway activation from organelle morphology requires that distinct pathways produce structurally distinguishable changes amenable to quantitative measurement. To evaluate whether Golgi architecture satisfies this prerequisite, we treated MRC-5 human lung fibroblasts with a panel of pharmacological compounds targeting defined signaling cascades: forskolin (10 µM, 30 min; cAMP/PKA), anisomycin (50 µg/mL, 60 min; JNK/p38 MAPK), TPA (100 nM, 60 min; PKC), and thapsigargin (2 µM, 60 min; ER stress/Ca²⁺). Each was selected for its well-characterized cascade and documented effects on Golgi organization ^16, 17^, with vehicle-treated cells (0.1% DMSO) serving as the baseline reference (**Table 1**). Fluorescence microscopy of GM130-immunolabeled cells revealed that these four pathway-active compounds generated visually distinguishable Golgi morphologies distributed along a fragmentation continuum (**Figure 1A**). Relative to the compact, crescent-shaped perinuclear ribbons of vehicle-treated cells, forskolin induced tighter cisternal packing and reduced ribbon extension, yielding a condensed Golgi profile consistent with PKA-mediated enhancement of anterograde transport and membrane consolidation ^16^, anisomycin produced partial ribbon dispersal with increased inter-cisternal spacing, reflecting JNK-dependent (and possibly p38-dependent) phosphorylation of the stacking protein GRASP65 ^32^; and TPA caused fragmentation into multiple discrete perinuclear elements, attributable to PKCα-mediated phosphorylation of GRASP55 that disrupts cisternal stacking ^16, 33^. Thapsigargin served a dual role, representing the ER stress/Ca²⁺ activation class while also functioning, given its reliable induction of Golgi fragmentation, as the positive control for the fragmented phenotype ^16^; as expected, it drove the most extensive fragmentation, dispersing the Golgi into ministacks throughout the cytoplasm. Notably, TPA and thapsigargin converge on the same proximal PKCα/GRASP55 axis but differ in their upstream trigger: TPA activates PKCα directly through phorbol-ester binding, whereas thapsigargin elevates cytosolic Ca²⁺ via SERCA inhibition, which in turn activates PKCα and drives GRASP55 phosphorylation ^17^. This Ca²⁺/PKCα-dependent fragmentation is thought to occur largely independently of the unfolded protein response, with the stronger, more sustained Ca²⁺-driven PKCα activation elicited by thapsigargin consistent with the more complete fragmentation observed, whereas the unfolded protein response develops in parallel and underlies its ER-stress transcriptional signature rather than necessarily serving as the sole proximal cause of Golgi disassembly ^17^. CHIR99021 (3 µM, 24 h; Wnt/β-catenin), was included as a specificity probe: although it potently activates canonical Wnt signaling through GSK-3β inhibition, no direct effect on Golgi morphology has been reported, and CHIR99021-treated cells indeed retained intact perinuclear ribbons indistinguishable from vehicle. The correspondence between each pathway and its observed Golgi phenotype is summarized schematically in **Figure 1B**.

**Figure 1.**
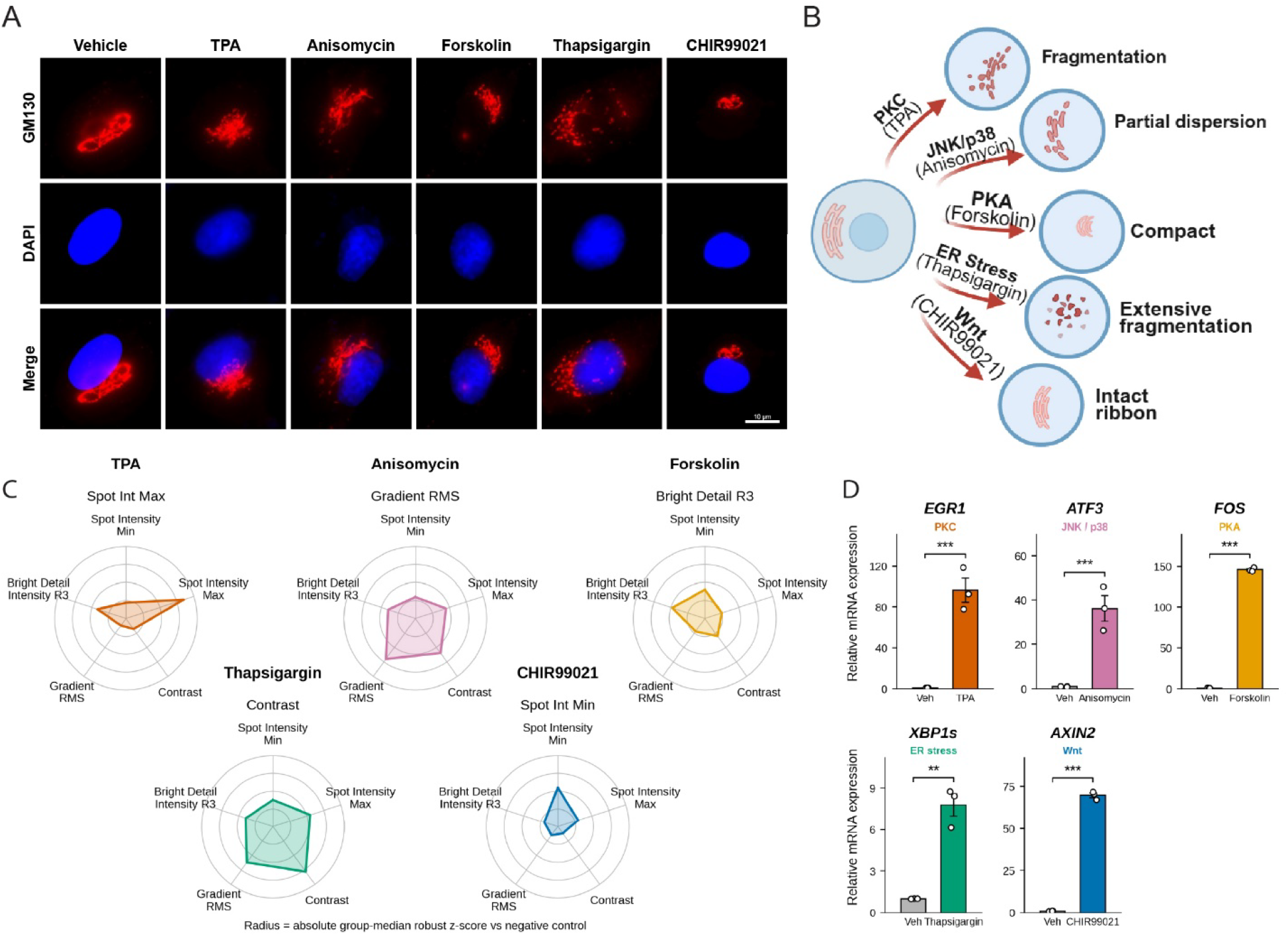
Signaling pathway activation produces distinct, quantifiable Golgi morphologies. **(A)** Representative fluorescence microscopy images of MRC-5 human lung fibroblasts immunolabeled for GM130 (cis-Golgi marker, red) and counterstained with DAPI (nuclei, blue) following treatment with the indicated pharmacological compounds. Vehicle-treated cells (0.1% DMSO) display compact perinuclear Golgi ribbons with crescent-shaped morphology. Forskolin (10 µM, 30 min) induces ribbon compaction with tighter cisternal packing and reduced ribbon extension, yielding a condensed perinuclear Golgi profile. CHIR99021 (3 µM, 24 h) retains intact perinuclear Golgi ribbons indistinguishable from vehicle, consistent with its role as a morphological negative control. Anisomycin (50 µg/mL, 60 min) causes partial ribbon dispersal with increased inter-cisternal spacing. TPA (100 nM, 60 min) causes fragmentation into multiple discrete Golgi elements distributed around the perinuclear region. Thapsigargin (2 µM, 60 min) induces extensive fragmentation into dispersed ministacks throughout the cytoplasm. Scale bar, 10 µm. **(B)** Schematic summarizing the correspondence between each signaling pathway and its observed Golgi phenotype along a compaction-to-fragmentation continuum: PKA (forskolin)-driven compaction, JNK/p38 (anisomycin)-associated partial dispersal, PKC (TPA)-driven fragmentation, and ER stress/Ca²⁺ (thapsigargin)-driven extensive fragmentation, with Wnt (CHIR99021) producing no direct structural change. **(C)** Morphometric signatures of pathway activation, shown as radar plots. For each compound, intensity-and texture-based descriptors computed from single-cell GM130 images using IDEAS software (Amnis/Cytek) were summarized as the absolute group-median robust z-score relative to the negative control, with each radar axis representing one descriptor (spot intensity min, spot intensity max, contrast, gradient RMS, and bright detail intensity R3) and the radial distance from the center encoding the magnitude of deviation from control (radius = absolute group-median robust z-score vs negative control). Each compound (TPA, anisomycin, forskolin, thapsigargin, CHIR99021) traces a distinct radar profile dominated by a different axis (panel title), recapitulating the morphological continuum: TPA by spot intensity max, anisomycin by gradient RMS, forskolin by bright detail intensity R3, and thapsigargin by contrast, whereas CHIR99021 traces a small, near-centered profile, indicating minimal deviation from control across all axes. **(D)** Pathway-specific transcriptional validation by qPCR. Relative mRNA expression of a canonical downstream target for each pathway in compound-treated versus vehicle (Veh) cells, EGR1 (PKC), ATF3 (JNK/p38), FOS (PKA), XBP1s (ER stress), and AXIN2 (Wnt). Bars show mean ± SEM (n = 4); Statistical significance was assessed by Welch’s two-sample *t*-test on log10 transformed fold-change values (ΔΔCt method), with normalization to GAPDH and vehicle control; *\*\*P* < 0.01, *\*\*\*P* < 0.001.

**Figure 2.**
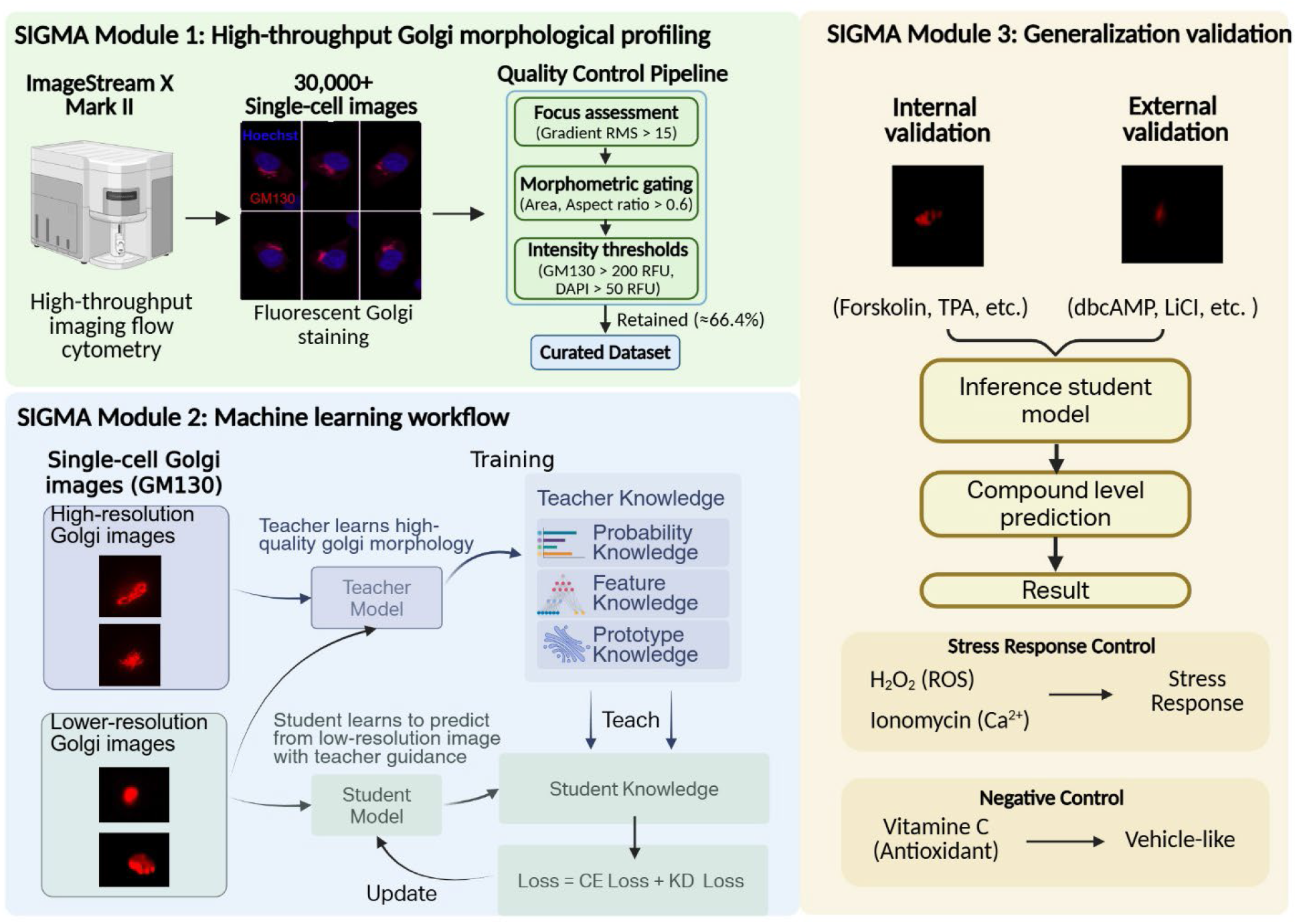
The SIGMA framework: a three-module strategy for morphology-based pathway classification. Schematic illustration of the three integrated modules comprising the SIGMA computational framework. **Module 1 (High-throughput morphological profiling):** MRC-5 fibroblasts are treated with pharmacological pathway activators, immunolabeled for GM130, and analyzed by ImageStreamX Mark II imaging flow cytometry at 60× magnification. Multi-stage quality control filtering (focus assessment, morphometric gating, fluorescence intensity thresholds) yields a quality-controlled dataset of single-cell Golgi images. Comprehensive single-cell morphological features are extracted, capturing signal intensity, texture, structural compactness, symmetry, and spot-level fragmentation. **Module 2 (Machine-learning workflow):** A teacher model learns high-quality Golgi morphology from high-resolution images and transfers three forms of teacher knowledge, soft pathway probabilities, feature embeddings, and class prototypes, to a student model trained on lower-resolution Golgi images. The student is optimized using supervised cross-entropy (CE) loss together with knowledge-distillation (KD) loss, producing the lower-resolution inference model. **Module 3 (Generalization testing):** The trained classifier is applied without retraining in two compound-level settings: internal testing on the reserved ImageStream test set and external testing on chemically distinct compounds, including pathway-matched activators, cross-pathway perturbations, and a negative control compound without expected Golgi effects. Successful compound assignment, especially for mechanistically related but structurally distinct compounds, tests whether classification reflects pathway biology.

**Table 1.** Pharmacological perturbation panel for signaling pathway activation.

| Compound | Concentration | Duration | Target<br>Pathway | Expected Morphological Effect |
| --- | --- | --- | --- | --- |
| Forskolin | 10 μM | 30 min | cAMP/PKA | Ribbon compaction <sup>15</sup> |
| CHIR99021 | 3 μM | 24 h | Wnt/β-<br>catenin | No direct Golgi-structural effect;<br>included as morphological negative<br>control (apparent separability at 24<br>h attributed to cell-cycle confound) |
| Anisomycin | 50 μg/mL | 60 min | JNK/p38<br>MAPK | Partial dispersal <sup>54</sup> |
| TPA | 100 nM | 60 min | PKC | Fragmentation via PKCα-mediated<br>GRASP55 phosphorylation <sup>16</sup> |
| Thapsigargin | 2 μM | 60 min | ER<br>stress/$\text{Ca}^{2+}$ | Fragmentation <sup>55</sup> (positive control) |
| Vehicle | 0.1% DMSO | 60 min | None | Baseline morphology |

While these qualitative observations demonstrated that pathway-specific morphologies exist, machine learning classification further requires that such differences be captured by quantitative features. To address this, we extracted a panel of intensity-and texture-based morphological descriptors from each GM130 labeled Golgi fluorescent image and, for every compound, summarized the magnitude of feature deviation from the negative control as the absolute group-median robust z-score **(Figure 1C)**. Across five representative descriptors, including spot intensity max, spot intensity min, contrast, gradient RMS, and bright detail intensity R3, each compound produced a distinct feature signature dominated by a different descriptor, recapitulating the qualitative continuum. TPA was dominated by spot intensity max, consistent with fragmentation into multiple bright, discrete Golgi elements; anisomycin by gradient RMS, consistent with the increased edge and texture heterogeneity of partial ribbon unlinking; forskolin by bright detail intensity R3, consistent with condensation of the ribbon into a compact perinuclear structure; and thapsigargin by contrast, consistent with extensive fragmentation and dispersal of ministacks throughout the cytoplasm. CHIR99021 produced only minimal deviation across all descriptors, indistinguishable in magnitude from the negative control, confirming the absence of pathway-driven Golgi remodeling and validating its role as a morphological negative control. Each treatment condition occupied a distinct region of this multivariate feature space, indicating that the extracted features carry sufficient discriminating information to support automated classification.

Having established that Golgi morphological differences are both pathway-specific and quantitatively measurable, we next verified that these changes reflect on-target signaling engagement rather than non-specific compound effects. qPCR analysis of a canonical downstream transcriptional target for each pathway confirmed the expected program: EGR1 (PKC; TPA), ATF3 (JNK/p38; anisomycin), FOS (PKA; forskolin), XBP1s (ER stress; thapsigargin), and AXIN2 (Wnt; CHIR99021), each significantly induced relative to vehicle **(Figure 1D)**. The concordance between Golgi morphological phenotypes and pathway-specific transcriptional markers indicates that the structural signatures are associated with the intended signaling perturbations.

These results satisfied the biological prerequisites for machine learning-based pathway classification. Golgi morphology encodes pathway-specific information along a quantifiable fragmentation continuum: from compaction under PKA activation, through partial dispersal under JNK/p38 stress signaling and fragmentation under PKC activation, to extensive fragmentation under ER stress/Ca^2+^ signaling, with each phenotype validated by independent biochemical and transcriptional evidence. The inclusion of CHIR99021, which activates Wnt signaling without altering Golgi structure, provides a built-in control for assessing classifier specificity against false-positive predictions. Having established this foundation using conventional microscopy, we next sought to scale the analysis to the large, statistically powered populations required for robust machine learning by transitioning to high-throughput imaging flow cytometry.

### The SIGMA framework: a three-module strategy for morphology-based pathway classification

The fluorescent microscopy analysis established that signaling pathway activation is associated with quantifiable, pathway-specific Golgi morphological changes **(Figure 1)**. Conventional microscopy is limited in throughput, typically analyzing dozens to hundreds of cells, which is insufficient for training robust machine learning classifiers ^34^. To address this scaling challenge while maintaining biological validity, we developed **SIGMA (SIgnaling via Golgi Morphology Analysis)**, a computational framework comprising three integrated modules **(Figure 2).** *Module 1* employs imaging flow cytometry to generate population-scale morphological profiles with rigorous quality control. Module 2 links these two image regimes by using teacher-guided distillation, allowing fine morphological cues learned from limited high-resolution Golgi crops to guide a student model trained on large-scale, lower-resolution ImageStream images. Module 3 tests generalization using chemically distinct compounds that engage the same signaling pathways, thereby testing whether learned representations reflect pathway biology.

### SIGMA Module 1: scaling Golgi morphological analysis with imaging flow cytometry

Training robust machine learning classifiers requires datasets substantially larger than those achievable through conventional microscopy. Imaging flow cytometry using the ImageStreamX Mark II platform resolves this scaling challenge by combining the statistical power of conventional cytometry (several thousand cells per condition) with subcellular spatial resolution sufficient for organelle morphology assessment **(Figure 3A)**. We selected GM130 as the structural reporter based on three considerations: its specific localization to *cis*-Golgi membranes through GRASP65 complexes provides reliable organelle labeling without cytoplasmic background; it remains largely membrane-associated under most of the perturbations used, supporting consistent visualization across treatment conditions; and GM130-based morphological assessment has been validated in prior imaging flow cytometry studies, enabling methodological comparison ^35^.

**Figure 3.**
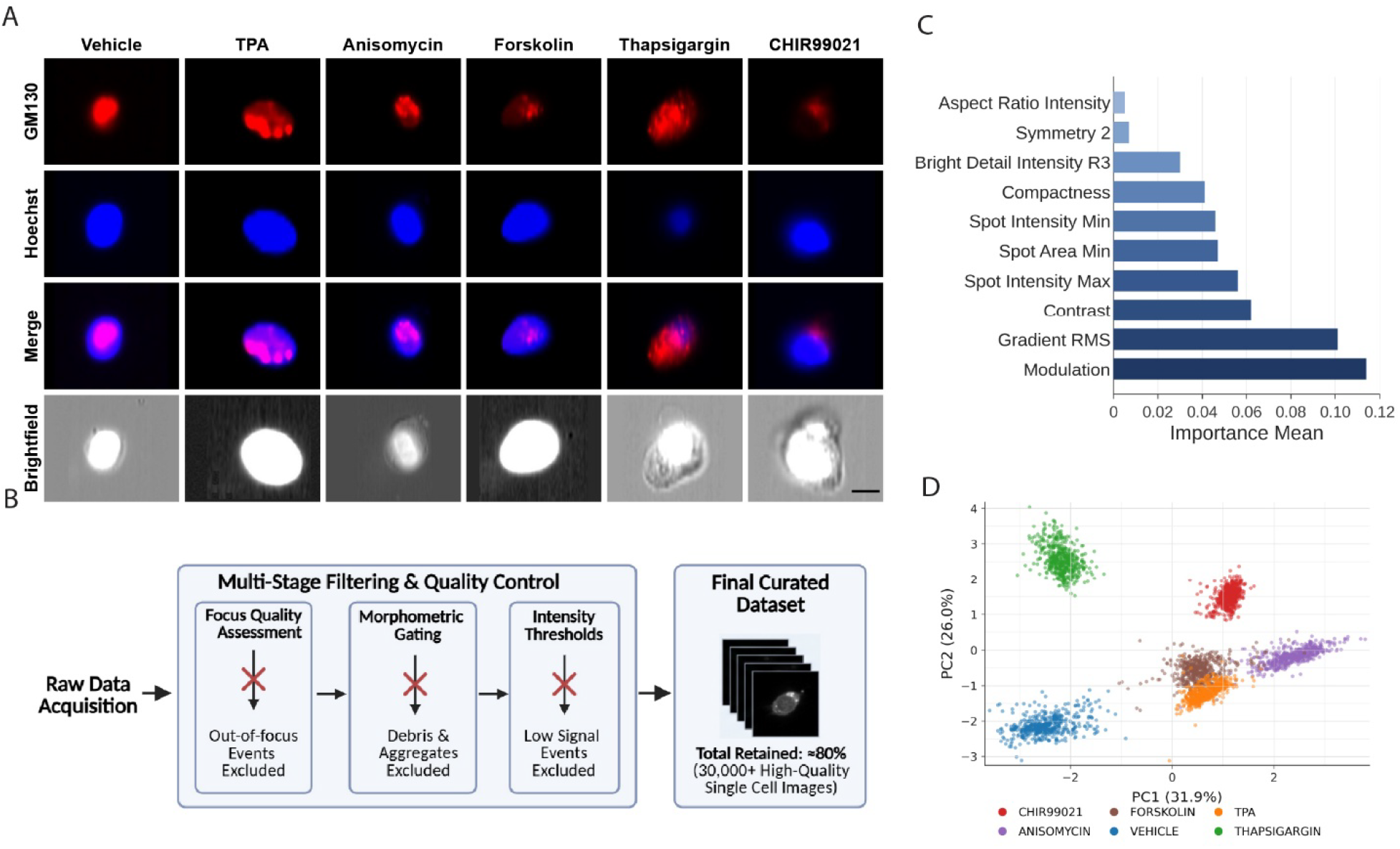
SIGMA Module 1: scaling Golgi morphological analysis with imaging flow cytometry. **(A)** Representative single-cell images acquired on the ImageStreamX Mark II platform at 60× magnification. For each treatment condition, four imaging channels are shown: brightfield (cell morphology), GM130 immunofluorescence (Golgi apparatus), Hoechst (nucleus), and merged composite. Images illustrate the subcellular spatial resolution achievable by imaging flow cytometry, sufficient for resolving Golgi architectural features including ribbon integrity, fragmentation state, and perinuclear positioning. Scale bar, 10 µm. **(B)** Multi-stage quality control pipeline for dataset curation. Raw imaging flow cytometry acquisitions are subjected to sequential filtering: (i) focus quality assessment using gradient root-mean-square (RMS) metrics on brightfield images (threshold: gradient RMS ≥ 15), removing out-of-focus events (∼8% of raw acquisitions); (ii) morphometric gating based on cell area and aspect ratio (Aspect Ratio_M01 ≥ 0.6), excluding cellular debris and multicellular aggregates (∼7% of remaining events); (iii) immunofluorescence intensity thresholds for GM130 (>200 RFU) and Hoechst (>50 RFU), ensuring adequate organelle and nuclear labeling (∼5% of otherwise acceptable events). Overall, 21,938 of 33,061 images were retained after quality control (66.4%). **(C)** Interpretability by permutation feature importance analysis of extracted Golgi morphological descriptors: Modulation emerged as the most influential descriptor, indicating that variation in Golgi signal heterogeneity captures a major component of pathway-relevant morphology, with spot-based and compactness features also contributing. This descriptor-level analysis provides correlative context and is distinct from the image-based distilled classifier used for prediction. **(D)** Principal component analysis (PCA) workflow for the sampled single-cell Golgi morphological feature profiles. From the quality-controlled ImageStream dataset, 3,000 single-cell profiles were sampled and represented by the extracted Golgi morphological descriptors, including intensity-, texture-, compactness-, symmetry-, and spot-level features. The descriptor matrix was standardized and projected onto the first two principal components; each point represents one sampled cell and is colored by treatment condition. PC1 and PC2 accounted for 31.9% and 26.0% of the variance, respectively (57.9% total).

Because machine learning classifiers can learn technical artifacts, we applied rigorous quality control before training, using a sequential four-stage gating strategy in IDEAS software in which each gate enriches for high-quality single-cell images **(Figure 3B)**. (1) Focus was assessed by brightfield Gradient RMS, a texture-based sharpness metric; events below a threshold of 15 were excluded as out-of-focus (∼8% of raw acquisitions). (2) Single cells were selected by brightfield Area versus Aspect Ratio, gating out doublets and aggregates (Aspect Ratio_M01 < 0.6; ∼7% of remaining events). (3) A Hoechst gate (>50 RFU) removed enucleated debris and apoptotic bodies, and (4) a GM130 gate (>200 RFU) removed cells with failed or weak labeling (together ∼5% of otherwise acceptable events). The resulting dataset spanned all six experimental conditions, with condition-specific retention rates ranging from 39.4% to 89.8% for a pooled retention of 66.4% **(Table 2)**.

**Table 2.** Dataset composition following quality control.

| **Condition** | **Raw Events** | **Post-QC Events** | **Retention (%)** |
| --- | --- | --- | --- |
| Vehicle | 3759 | 1,480 | 39.4 |
| Forskolin | 2844 | 1,653 | 58.1 |
| CHIR99021 | 7146 | 3,912 | 54.7 |
| Anisomycin | 4107 | 3,419 | 83.2 |
| TPA | 7873 | 7,070 | 89.8 |
| Thapsigargin | 7332 | 4,404 | 60.1 |

To characterize Golgi morphology, we extracted multiple quantitative descriptors from each single-cell image using the built-in tools in IDEAS^®^. Intensity and texture features (modulation, gradient RMS, contrast, bright detail intensity R3, aspect ratio intensity) captured signal heterogeneity and edge detail; spot-level features (spot intensity max, spot intensity min, spot area min) captured the brightness and size of fragmented Golgi regions; and compactness and symmetry quantified overall geometric organization. Together these descriptors summarized Golgi dispersion, fragmentation, intensity heterogeneity, and structural regularity.

To examine whether the extracted morphological descriptors captured treatment-dependent structure, we used permutation feature importance to quantify the contribution of individual descriptors **(Figure 3C)** and principal component analysis (PCA) to visualize treatment-associated variation in single-cell morphology **(Figure 3D)**. The projection organized cells by treatment conditions, with several groups occupying distinct regions and forskolin and TPA showing adjacent, partially overlapping distributions, consistent with a graded continuum of Golgi morphological states. Together, these patterns indicate that the extracted descriptors capture treatment-associated Golgi morphological variation while preserving biologically plausible overlaps among related cellular states, supporting their use as quantitative inputs for downstream pathway classification.

### SIGMA Module 2: pathway classification by teacher-student knowledge distillation

The preceding sections established two requirements for morphology-based pathway inference: signaling activation produces measurable Golgi phenotypes **(Figure 1)**, and imaging flow cytometry can scale those phenotypes to population-level single-cell datasets **(Table 2 and Figure 3)**. Module 2 was designed around the different roles of these image sources. Although both platforms acquire at 60× magnification, magnification and resolution are distinct: the ImageStream’s single-plane, in-flow acquisition and coarser pixel sampling give it lower effective spatial resolution than the coverslip microscope, so the fine Golgi substructure resolved in the microscopy crops (individual cisternae and dispersed ministacks) is only partially captured in the ImageStream images. High-resolution microscopy preserves fine Golgi structure, but high-resolution single-cell Golgi crops are limited in number because they require labor-intensive acquisition, quality review, and manual cropping. In contrast, ImageStream-derived lower-resolution images can be acquired at much larger scale with substantially less manual processing, making them the realistic input for screening-scale classification, even though their lower resolution can obscure subtle structural cues needed to distinguish pathway-specific Golgi states. Training only a high-resolution classifier would therefore learn from the most informative images but would not directly solve the deployment problem, because routine inference must operate on lower-resolution ImageStream images. We therefore framed Module 2 as teacher-guided transfer: using scarce, information-rich high-resolution crops to define morphology-sensitive guidance while training the deployed classifier directly on abundant lower-resolution images and pathway labels.

For this classification task, SIGMA used a morphology-aware teacher-student distillation framework. The teacher model was first trained on 139 manually curated high-resolution single-cell Golgi crops and then frozen as a reference representation of fine Golgi morphology. During distillation, each lower-resolution ImageStream training image was passed through both the frozen teacher and the student, so teacher outputs could guide the student without requiring one-to-one matched high-resolution and lower-resolution image pairs. The student model was trained on normalized lower-resolution ImageStream GM130/Golgi images and received three complementary supervision signals from the teacher **(Figure 4A)**: probability knowledge, which transfers class-level soft pathway relationships from teacher output distributions; feature knowledge, which aligns the student embedding with the teacher’s morphology-sensitive representation; and prototype knowledge, which anchors student features to class-level morphology prototypes computed from the high-resolution crops ^29, 36, 37, 38^.

**Figure 4.**
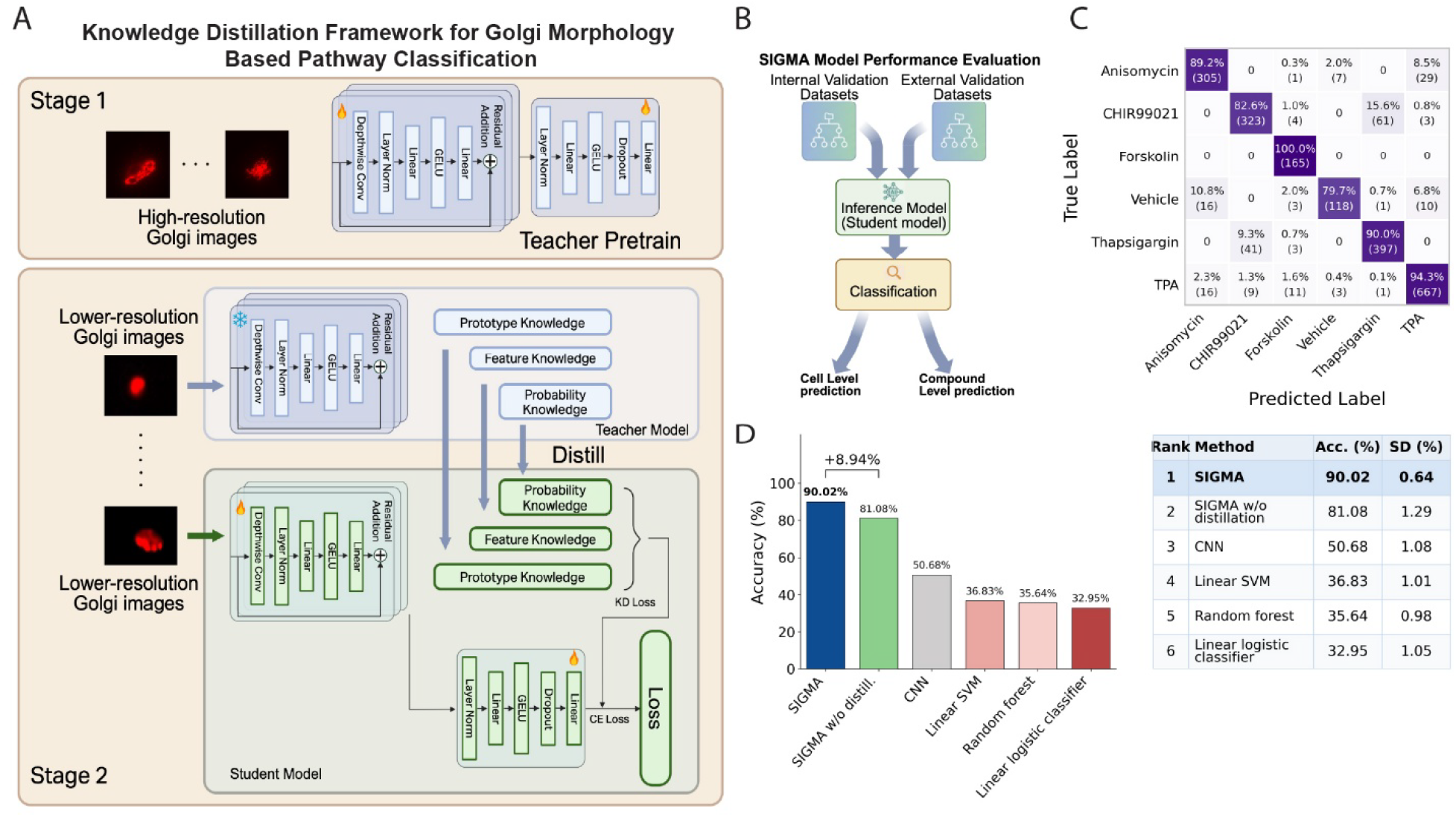
SIGMA Module 2: pathway classification by teacher–student knowledge distillation. **(A)** Two-stage knowledge-distillation framework for Golgi morphology-based pathway classification. In Stage 1, a teacher network is pretrained on high-resolution Golgi images and then frozen. In Stage 2, lower-resolution Golgi images are processed by the frozen teacher and the trainable student. Prototype, feature, and probability knowledge from the teacher are transferred to the student through a composite knowledge-distillation (KD) loss, while the student classification head is jointly optimized with supervised cross-entropy (CE) loss. After training, inference is performed entirely by the lower-resolution student across six labels. **(B)** Validation workflow. Validation of the trained student model was performed at both the cell and compound levels. In Module 2, each single-cell image was independently classified to generate a predicted label. In Module 3, a group of single-cell images from the same compound was provided to the model to generate a single compound-level label (Figure 5). **(C)** Row-normalized confusion matrix on the cell-level test set. Rows are true pathway labels and columns are predicted labels; each cell shows the count and row-normalized percentage, and color intensity encodes the row-normalized percentage (0–100%). Overall cell-level test accuracy was 90.02%. Diagonal values denote per-class recall: anisomycin 89.2%, CHIR99021 82.6%, forskolin 100.0%, vehicle 79.7%, thapsigargin 90.0%, and TPA 94.3%. Off-diagonal entries reflect genuine morphological similarity among certain signaling states. **(D)** Comparison of classification accuracy across methods on the same cell-level test set.A bar chart shows cell-level test accuracy for each method, and the accompanying table reports the corresponding accuracy (Acc.) and standard deviation (SD). SIGMA achieved the highest accuracy (90.02% ± 0.64%), exceeding the SIGMA w/o distillation (81.08% ± 1.29%), a CNN trained from scratch (50.68% ± 1.08%), a linear SVM (36.83% ± 1.01%), a random forest (35.64% ± 0.98%), and a linear logistic classifier (32.95% ± 1.05%). The 8.94 percentage-point gain over the SIGMA w/o distillation indicates that teacher-guided representation transfer improves lower-resolution classification beyond non-distilled baselines.

These three distillation signals were included to transfer complementary forms of high-resolution morphological information into the lower-resolution student. Probability knowledge provides class-level soft supervision, allowing the student to inherit the teacher’s relative pathway relationships. Feature knowledge provides instance-level morphology alignment, encouraging each lower-resolution representation to preserve structural cues captured by the teacher. Prototype knowledge provides class-level morphological anchors, guiding lower-resolution features toward high-resolution class prototypes and stabilizing discrimination among visually similar pathway states. Together, these signals allow SIGMA to use high-resolution images as an information source during teacher training and prototype construction while keeping the final classifier optimized for lower-resolution ImageStream inputs at inference.

For probability knowledge, teacher and student logits were temperature-scaled and compared as output distributions:

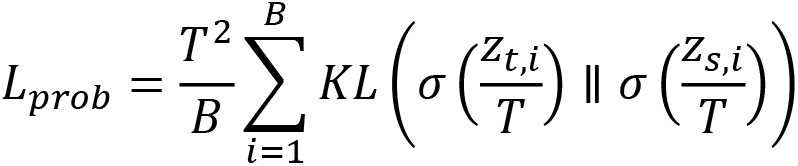

where σ(.) denotes softmax, KL denotes Kullback-Leibler divergence, zₜ and zₛ are the teacher and student logits, and T = 2.0 was used in training.

For feature knowledge, rᵢ denotes the L2-normalized projected student feature and tᵢ denotes the L2-normalized teacher feature generated by the same lower-resolution image through the frozen teacher. Feature alignment was optimized using cosine distance ^36^:

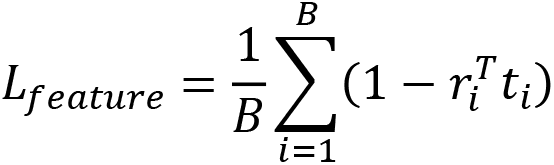

This term encouraged the lower-resolution student encoder to approximate the morphology-sensitive representation produced by the teacher.

For prototype knowledge, class prototypes were computed from the high-resolution teacher crops ^37^. For each class c, the prototype was the normalized mean of normalized teacher features from high-resolution crops belonging to that class:

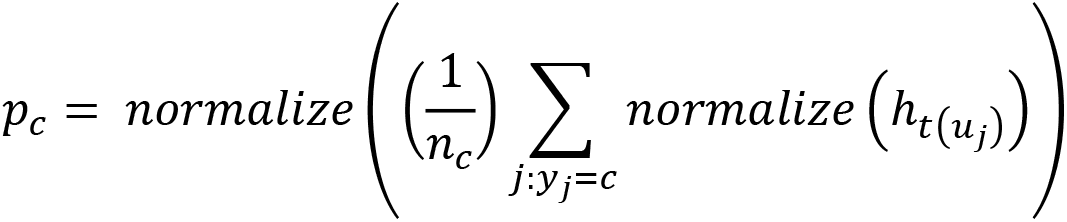

where uⱼ denotes a high-resolution teacher crop and n(c) is the number of teacher crops in class c. During student training, the projected student feature rᵢ was compared with all class prototypes, and the prototype loss was computed as cross-entropy over prototype-similarity logits:

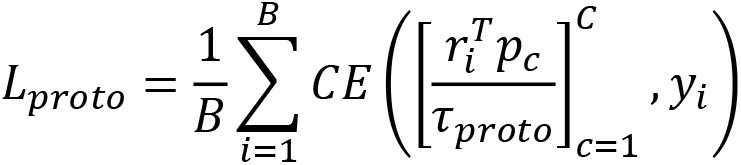

where τ_ₚᵣₒₜₒ_ is the prototype temperature and yᵢ is the ground-truth label, and τₚᵣₒₜₒ = 0.2 was used in training.

The composite knowledge-distillation term was defined as the sum of the three teacher-guided distillation losses:

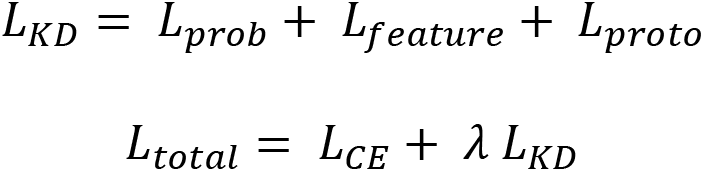

The λ balances the contributions of the loss terms, and λ was scheduled with a cosine-ramp schedule in training.

This design provided three practical advantages. First, the high-resolution teacher supplied a fine-morphology prior from the small curated crop set, allowing structural information that may be weak or partially unresolved in ImageStream images to influence student training. Second, because the student was still trained directly on lower-resolution ImageStream images and pathway labels, the deployed model learned discriminative features in the same image domain used at inference rather than relying on a high-resolution model that may not transfer cleanly to lower-resolution inputs. Third, after distillation, only the lower-resolution student was required for inference, reducing dependence on labor-intensive high-resolution acquisition and manual crop preparation. In this way, SIGMA used the high-resolution images where they were most valuable, as guidance for representation learning, while preserving the scalability of lower-resolution imaging for routine classification.

To evaluate the classification performance, we performed two complementary validation schemes: cell-level validation and compound-level validation **(Figure 4B).** In the cell-level validation, each single-cell image was independently classified, and accuracy was calculated across individual cells. In compound-level validation, predictions from all cells under the same treatment condition were aggregated to generate one compound-level prediction. The row-normalized confusion matrix showed strong diagonal dominance across all six labels **(Figure 4C)**, indicating that the classifier learned pathway-associated Golgi signatures. We also benchmarked SIGMA against the SIGMA w/o distillation and four conventional baselines on the same test set **(Figure 4D)**. SIGMA achieved the highest accuracy (90.02%), exceeding the SIGMA w/o distillation (81.08%), a CNN trained from scratch (50.68%), linear SVM (36.83%), random forest (35.64%), and linear logistic classifier (32.95%). These comparisons support Module 2: pathway information in Golgi morphology is present in the lower-resolution dataset, but teacher-guided deep feature learning extracts it more effectively than shallow classifiers, a scratch-trained CNN, or the same student trained without high-resolution guidance ^34, 38^.

### SIGMA Module 3: testing generalization to chemically distinct compounds

Having demonstrated cell-level test performance in Module 2, we next evaluated compound-level assignment in Module 3. A classifier that memorizes compound-specific features would fail when presented with chemically distinct molecules engaging the same signaling pathway. Conversely, successful classification of Module 3 test compounds based solely on Golgi morphological similarity to training data would provide evidence that SIGMA has learned generalizable pathway-associated features. Module 3 separated internal and external validation: internal validation used the reserved 10% held-out ImageStream test partition from the original compound/control set, whereas external validation used a new independent panel of chemically distinct compounds. In both settings, images were processed through the same imaging, quality-control, and lower-resolution image-normalization workflow used for student inference. The classification step applied the same frozen distilled student and inference procedure used for Module 2 validation **(Figure 4B)**. Cell-level class probabilities were aggregated within each compound or control condition to generate one compound-level label, and class assignments were generated without any model retraining **(Figure 5A).**

**Figure 5.**
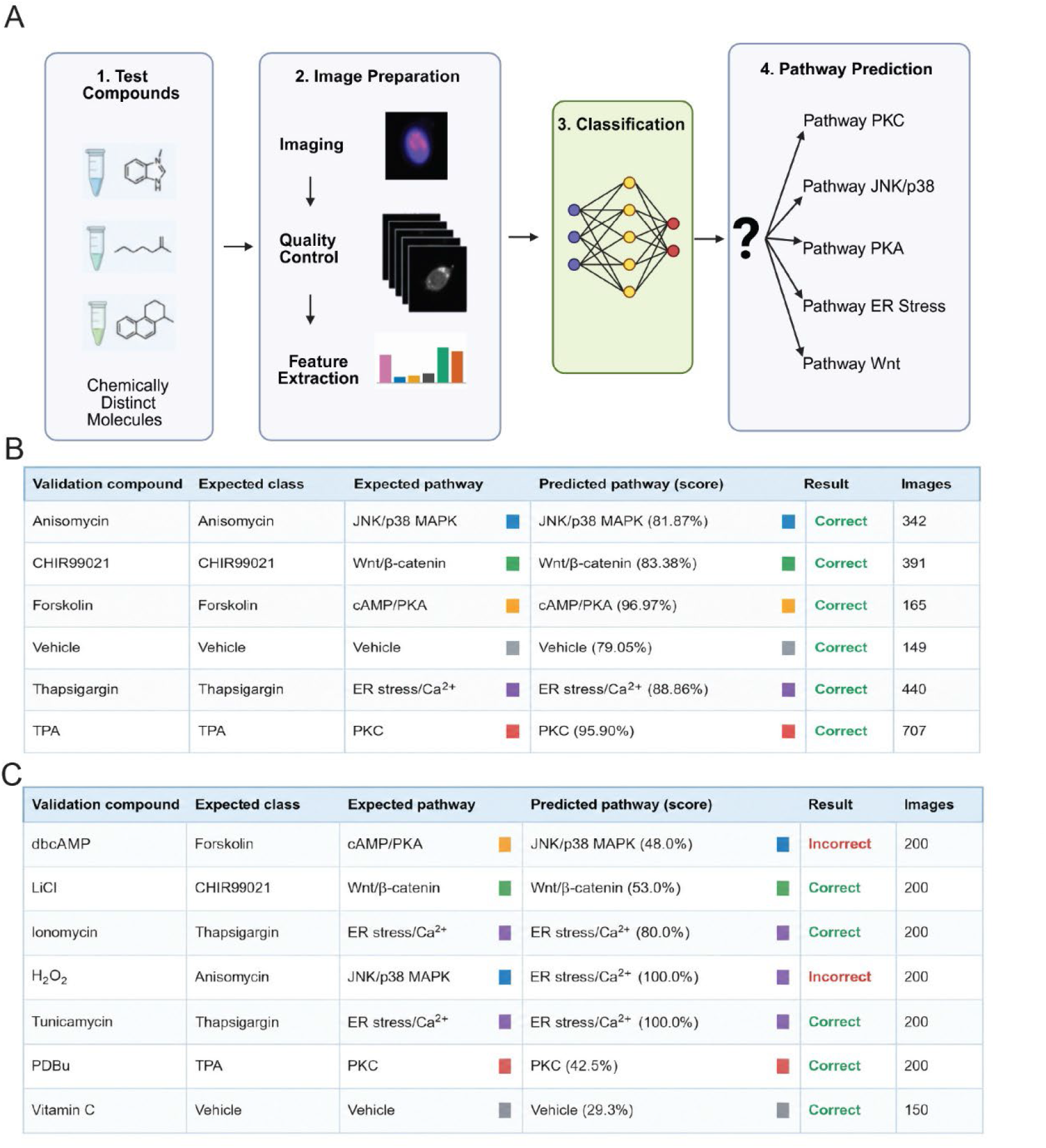
SIGMA Module 3: internal and external compound-level testing. **(A)** Schematic of the SIGMA generalization (prediction) workflow. The classifier trained in Module 2 is applied to chemically distinct test compounds without retraining: (1) novel compounds are treated; (2) cells are processed through the same image-preparation pipeline used for student inference, including imaging, quality-control gating, and lower-resolution image normalization; (3) the SIGMA model classifies each cell; and (4) a pathway prediction is assigned among the learned signaling classes, the question mark denoting the unknown assignment for a previously unseen compound. Correct prediction of mechanistically related but chemically distinct compounds tests whether classification reflects pathway biology. **(B)** Internal compound-level testing on the reserved test set. For each test condition or control, the table lists the expected class and pathway, the predicted pathway with the aggregated compound score, the classification result, and the number of single-cell images. All six labels were correctly assigned: anisomycin-JNK/p38 MAPK (81.87%), CHIR99021-Wnt/β-catenin (83.38%), forskolin-cAMP/PKA (96.97%), vehicle (79.05%), thapsigargin-ER stress/Ca²⁺ (88.86%), and TPA-PKC (95.90%), yielding 6/6 correct assignments. **(C)** External compound-level test panel of seven chemically distinct compounds evaluated without model retraining. The table lists the test compound, expected training class and pathway, SIGMA-predicted pathway with score, classification result, and number of images. Correct assignments were obtained for LiCl-Wnt/β-catenin (53.0%), ionomycin-ER stress/Ca²⁺ (80.0%), tunicamycin– ER stress/Ca²⁺ (100.0%), PDBu-PKC (42.5%), and vitamin C-vehicle (29.3%). dbcAMP was assigned to JNK/p38 MAPK (48.0%) instead of cAMP/PKA, and H₂O₂ was assigned to ER stress/Ca²⁺ (100.0%) instead of JNK/p38 MAPK, yielding five of seven correct compound assignments.

We assembled an external test panel of seven chemically distinct compounds to test whether the classifier generalizes from its training compounds to chemically unrelated molecules engaging the same signaling pathways. Six pathway-matched compounds were each expected to map to the training class engaged by the same cascade: dibutyryl-cAMP (dbcAMP; cAMP/PKA), LiCl (Wnt/β-catenin), the Ca²⁺ ionophore ionomycin and the N-glycosylation inhibitor tunicamycin (ER stress/Ca²⁺), the oxidative-stress agent H₂O₂ (JNK/p38 MAPK), and the phorbol ester PDBu (PKC); the antioxidant vitamin C was included as a negative control expected to remain at the basal class. For each compound, cell-level class probabilities were aggregated to a single class assignment, and a prediction was scored as correct when the top-ranked class matched the expected pathway (or, for vitamin C, the vehicle/control class).

Internal compound-level testing was performed first on the reserved test set: after aggregating cell-level class probabilities within each original compound/control label, all six internal labels were assigned to their expected class (6/6, 100%) **(Figure 5B)**. We then applied the trained SIGMA classifier to the external seven-compound test panel and evaluated predictions at the compound level **(Figure 5C)**. For the external test panel, five of seven compounds were assigned to their expected pathway or control category, corresponding to 71.4% external compound-level accuracy. Pathway-matched compounds engaging PKC (PDBu) and Wnt/β-catenin (LiCl), the ER stress/Ca²⁺ agents ionomycin and tunicamycin, and the vitamin C negative control were all classified to their expected class, indicating that SIGMA captured pathway-associated morphology but not compound identity alone; ionomycin and tunicamycin engage the ER stress/Ca²⁺ axis through Ca²⁺ influx and N-glycosylation blockade, respectively, mechanisms distinct from the SERCA inhibition used by the training compound thapsigargin. The two misclassifications both involved the stress-responsive classes: the PKA-class compound dbcAMP was assigned to the JNK/p38 (anisomycin) class, and the oxidative-stress agent H₂O₂, expected to engage JNK/p38, was assigned instead to the ER stress/Ca²⁺ class. These errors indicate that the JNK/p38 MAPK and ER stress/Ca²⁺ fragmentation phenotypes overlap in morphological feature space, marking a boundary where morphology-only classification may require additional molecular context.

## Discussion

Our observation that Golgi morphology carries pathway-specific information places the Golgi within a broader view of cellular architecture as a readout of molecular functional states. The central advance of this work is to establish organelle morphology as a multiplexed pathway fingerprint: SIGMA discriminates among multiple discrete signaling states from structure alone, extending morphofunction from the magnitude of one function to the identity of the active pathway. This principle has recent empirical support. Ding *et al.* showed that mitochondrial shape predicts metabolic activity^6^, while De Vries et al. and Longden *et al.* showed that whole-cell morphological profiles predict signaling-pathway perturbations and drug-resistance mechanisms complementary to molecular markers ^7, 8^. Our findings extend this principle to the Golgi. Its broad regulatory connectivity may make it well suited as a morphological integrator: unlike organelles with more circumscribed functions, it receives inputs from cell-cycle machinery, stress-response pathways, secretory-demand signals, and polarity cues ^39, 40, 41, 42, 43^. To the extent that each cascade produces a distinguishable architectural state, these states constitute the morphological signature SIGMA is trained to classify.

Teacher-guided knowledge distillation enabled SIGMA to work with high-throughput imaging. At the cell level, the distilled student reached 90.02% test accuracy and a macro-averaged F1 score of 89.33%, compared with 81.08% for the non-distilled student, indicating that teacher guidance improved classification from lower-resolution ImageStream images. The distilled student also outperformed a scratch-trained CNN (50.68%) and conventional machine-learning baselines, including linear SVM (36.83%), random forest (35.64%), and linear logistic classifier (32.95%). These comparisons support the use of distillation. This design reflects the experimental reality: high-resolution Golgi crops preserve fine structural information but are limited and labor-intensive to acquire, while lower-resolution ImageStream images are abundant and scalable but may underrepresent subtle morphological cues. By using the high-resolution teacher as a source of morphology-sensitive supervision during training, SIGMA transfers fine structural information into a student model that is trained and deployed on lower-resolution images. After distillation, only the lower-resolution student is required for inference, reducing dependence on large-scale high-resolution acquisition while preserving sensitivity to pathway-associated Golgi morphology. In this proof-of-concept, distillation allowed a model trained partly on high-content images to operate on screening-scale data without a measured loss of pathway sensitivity.

SIGMA complements molecular pathway assays. Whereas targeted assays measure specific components, morphological classification provides an integrative readout capturing convergent effects of signaling on organelle architecture, which may be advantageous where activation produces distributed effects not captured by single-marker assays. To characterize the morphology underlying these classes, Permutation feature importance analysis was applied to extracted descriptors, providing correlative explanation. It identified Modulation, a measure of Golgi signal heterogeneity, as the most influential descriptor, with spot-based and compactness features also contributing (Figure 3C), mirroring the compaction-to-fragmentation continuum seen in the representative images.

Several limitations remain. The study used a single cell type, so generalization across lineages with different baseline Golgi architectures is undetermined and may require retraining. We examined individual pathway activation under controlled conditions, whereas physiological contexts involve combinatorial activation and crosstalk that may yield unrepresented states. Treatment durations also varied, from 30 min to 24 h. However, four of five activators and the vehicle shared short durations, so discrimination among these classes is unlikely to be driven by culture time. Finally, generalization to chemically distinct compounds was only partially established: the panel was small, and among the three chemically distinct pathway-matched activators, only PDBu was both correctly and specifically classified, yet PDBu shares the phorbol-ester scaffold of the training compound TPA and therefore serves as a same-class positive control rather than a fully independent structural test. Larger, chemically diverse panels will be needed to establish the scope of pathway-level generalization.

## Conclusions

This study demonstrates that Golgi morphology carries pathway-level information that deep learning can read: a teacher-student distillation strategy transfers representations learned from high-resolution images to a lightweight classifier that operates at the throughput of imaging flow cytometry, decoupling the resolution needed for feature learning from that needed at deployment. Correct classification of chemically distinct compounds absent from training indicates that the learned features capture pathway-associated morphology, though this generalization was compound-dependent. Together, these results establish morphological pathway fingerprinting as an interpretable, throughput-compatible readout of signaling state and a foundation for target-engagement screening and high-content phenotyping.

## Materials and Methods

### Cell culture and pharmacological treatments

Cell culture and pharmacological treatments were performed as previously described ^44, 45^. MRC-5 human lung fibroblast was obtained from ATCC. Cells were cultured in Dulbecco’s Modified Eagle Medium (DMEM) supplemented with 10% fetal bovine serum (FBS), 1% penicillin-streptomycin. The culture conditions were 37 °C and 5% CO2. Cells were passaged at 70-80% confluence. After gentle washing with Ca^2+^/Mg^2+^-free PBS, cells were digested with 0.05% trypsin-0.53 mM EDTA, and the process was terminated with an equal volume of complete DMEM. For signaling pathway analysis, cells were grown to 70-80% confluence and treated with specific agonists or inhibitors as follows: To activate PKC signaling, cells were treated with TPA (12-O-tetradecanoylphorbol-13-acetate) at 100 nM for 60 min. For JNK/p38 MAPK activation, cells were incubated with anisomycin (50 μg/mL) for 60 min. Wnt signaling was modulated by treating cells with the GSK-3β inhibitor CHIR99021 at 3 μM for 24 h. To induce cytosolic calcium release and ER/Golgi stress, cells were treated with thapsigargin (2 μM) for 60 min. To raise cAMP and activate PKA, adenylyl cyclase was directly stimulated with forskolin at 10 µM for 30 min. Cells were treated with ionomycin (1 μM), tunicamycin (5 μg/mL), dbcAMP (100 μM), LiCl (100 μM), H2O2 (25 μM), or vitamin C (50 µM) for 60 min, and PDBu (50 nM) for 30 min. DMSO (0.1%) was used as the vehicle control for all drug treatments. Pharmacological compounds were prepared as concentrated stock solutions in DMSO (forskolin, CHIR99021, anisomycin, TPA, thapsigargin, PDBu) or aqueous buffer (dbcAMP, LiCl, vitamin C) and diluted to working concentrations in complete medium immediately before treatment. Final DMSO concentration did not exceed 0.1% for any condition. Vehicle controls received 0.1% DMSO. A short-duration vehicle (60 min) was matched to the 30-60 min activators, and a duration-matched 24 h vehicle (0.1% DMSO, 24 h) was included as the reference for the 24 h CHIR99021 arm to isolate culture-time effects from pathway-specific remodeling

### Immunofluorescence staining and fluorescent microscopy

Immunofluorescence staining and fluorescent microscopy were performed as previously described ^46, 47, 48^. For immunofluorescence, MRC-5 cells were seeded on glass coverslips (6-well) and grown to ∼50-70% confluence. Cells were rinsed in PBS, fixed in 4% paraformaldehyde/PBS (15 min, on ice), washed (PBS, 3×), permeabilized with 0.1% Triton X-100/PBS (5 min), and blocked in 5% BSA/PBS (30 min, RT). Samples were incubated with anti-GM130 primary antibody (Cell Signaling, 1:150 in 5% BSA/PBS; 50 μL/coverslip) overnight at 4 °C, washed, and then with species-appropriate Alexa Fluor secondary antibody (1:1000, 30 min, RT, protected from light). Nuclei were counterstained with DAPI (1 μg/mL in PBS, 10 min), followed by two PBS washes and mounting in anti-fade medium. Images were acquired on an Echo Revolve fluorescence microscope (Discover Echo) using a 60× objective.

### RNA isolation and RT-qPCR

RNA isolation and RT-qPCR were performed as previously described ^49, 50, 51^. Total RNA was extracted from cultured MRC-5 cells using the Monarch Total RNA Miniprep Kit (New England Biolabs) per the manufacturer’s instructions, and purity and integrity were confirmed by A260/A280 ratio (>1.8). One microgram of total RNA was reverse-transcribed with the LunaScript RT SuperMix Kit (New England Biolabs) using random hexamer priming. Quantitative PCR was performed with Luna Universal qPCR Master Mix (New England Biolabs) on a CFX96 Real-Time PCR Detection System (Bio-Rad Laboratories) using transcript-specific primers, with amplification specificity confirmed by melt-curve analysis. GAPDH served as the internal reference gene, and relative mRNA expression was calculated using the 2^−ΔΔCt method and expressed as fold change relative to the indicated control group.

### Imaging flow cytometry

Imaging flow cytometry was performed as previously described ^52, 53^. Briefly, adherent MRC-5 cultures were gently dissociated with 0.05% trypsin-EDTA, neutralized in DMEM + 10% FBS, pelleted (300 × g, 5 min), fixed in 4% paraformaldehyde/PBS (10 min, RT), permeabilized with 0.1% Triton X-100/PBS (10 min) and blocked in 3% BSA/PBS (30 min). Cells were stained in suspension with anti-GM130 (1:100 in 5% BSA/PBS; 30 min RT), washed, incubated with species-appropriate Alexa Fluor secondary antibody (1:1000, 30 min, protected from light), counterstained with Hoechst (1 μg/mL, 10 min), and resuspended in PBS/5% BSA. Data were acquired on an ImageStreamX Mark II (Luminex/Cytek) at UAB flow cytometry core facility, using the 60× objective at low flow rate, with 405-nm excitation for Hoechst and 561-nm excitation matched to the Alexa Fluor 555 secondary fluorophore, brightfield was collected concurrently. For each condition, several thousand single-cell events were recorded. Analysis in IDEAS software proceeded by sequential gating of focused cells, singlets, and Hoechst-positive nuclei. Saturated images and debris were excluded.

### Data filtering

Before model training, lower-resolution ImageStream-derived Golgi images were filtered to remove insufficient-signal and artifact-containing samples. Images flagged as extreme dark, overexposed, low-information spot, black-line artifact, compact smooth blob, or lamp/bright-blob artifact were excluded. Retained images were normalized to 8-bit intensity values, organized by pathway label, and used as the model-ready lower-resolution dataset for student training and evaluation.

### Dataset and problem formulation

After image quality control, artifact removal, and lower-resolution image normalization, the model-ready lower-resolution dataset contained 21,938 single-cell GM130/Golgi images assigned to six labels. Images were stored as normalized 8-bit grayscale inputs and resized to 224 × 224 pixels before model input. The dataset was stratified by class into mutually exclusive training, cell-level test, and compound-level test sets using an 80%/10%/10% split, yielding 17,550 training images, 2,194 cell-level test images, and 2,194 compound-level test images. The training set was used to fit the student model, the cell-level test set was used for Module 2 performance evaluation and baseline comparison, and the compound-level test set was reserved for Module 3 internal testing.

The pathway-classification dataset was defined as D = {(xᵢ, yᵢ)} for i = 1,…,N, where xᵢ denotes the i-th single-cell lower-resolution Golgi image, yᵢ denotes its pathway label, N = 21,938, and C = 6. The classification task was to learn a model f that maps each input image xᵢ to one of the six pathway labels.

The high-resolution teacher dataset contained 139 manually curated single-cell Golgi crops across the same biological labels: anisomycin (n = 20), CHIR99021 (n = 20), forskolin (n = 26), vehicle (n = 29), thapsigargin (n = 20), and TPA (n = 24). These high-resolution crops were used only to train the teacher model and to compute class-level teacher prototypes.

### Model architecture

SIGMA used a teacher-student convolutional distillation framework. The teacher was a ConvNeXt-like classifier ^38^ trained on the manually curated high-resolution Golgi crops and then frozen. Because the high-resolution teacher dataset was limited in size (n = 139), no separate validation subset was withheld. All curated high-resolution crops were used to train the teacher. The teacher was subsequently frozen and was not evaluated as a standalone classifier; all reported classification performance was assessed using the independently held-out lower-resolution student test sets. During distillation, lower-resolution ImageStream GM130/Golgi images were passed through both the frozen teacher and the student to compute teacher-guided probability and feature targets, without requiring paired high-resolution and lower-resolution images. The student was another ConvNeXt-like classifier trained on lower-resolution ImageStream GM130/Golgi images and used for all inference.

The supervised classification term was computed over each mini-batch using cross-entropy between the student prediction and the ground-truth pathway label:

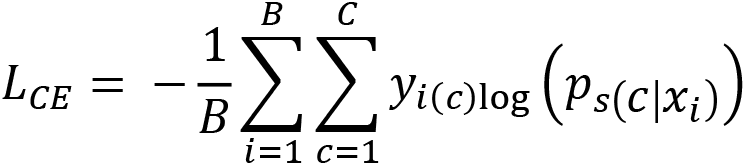

where B is the mini-batch size, c indexes pathway classes, y is the one-hot class indicator, and pₛ denotes the student class-probability output.

The student was trained for up to 200 epochs using AdamW optimization (learning rate 2 × 10⁻⁴, weight decay 5 × 10⁻⁴), label smoothing of 0.03, batch size 16, and a polynomial learning-rate decay.

Permutation feature importance analysis and PCA analysis were applied to extracted numeric Golgi morphological descriptors, with these descriptors, spanning intensity-, texture-, compactness-, symmetry-, and spot-level measures, providing feature-level context separate from the image-based distilled classifier.

### Baseline classifiers and comparison

To benchmark SIGMA against other approaches, we compared a linear SVM, linear logistic regression, random forest, and a CNN trained on the same six-class prediction task. All comparison methods used the same normalized lower-resolution ImageStream Golgi images and identical training and cell-level test partitions as SIGMA. The CNN directly processed the two-dimensional images, whereas the ImageStream images were flattened into one-dimensional vectors for input to the linear SVM, logistic regression, and random forest classifiers. Results are reported as the accuracy calculated on the cell-level test set ± the bootstrap-estimated SD.

### Statistics and Reproducibility

Image-based analyses used the acquisition, quality-control, artifact-removal, and intensity-filtering workflow described above. Raw and post-quality-control single-cell image counts for each treatment condition are reported in Table 2. No additional images were excluded after these pre-specified filtering steps. No formal power calculation was used to predetermine sample size.

For Module 2 model development, retained lower-resolution ImageStream-derived GM130/Golgi images were normalized and stratified by class into non-overlapping training, cell-level test, and compound-level test sets at the single-cell image level (80%/10%/10%; 17,550 training images, 2,194 cell-level test images, and 2,194 compound-level test images). training set was used for model fitting, and Module 2 cell-level performance evaluation and baseline comparisons were evaluated on the cell-level test set. Hyperparameters were selected using a stratified validation subset drawn exclusively from the training partition. After hyperparameter selection, the configuration was fixed and evaluated once on the held-out cell-level and compound-level test sets. Module 2 accuracy was computed at the cell level, with each image scored against its known label and reported together with the full error structure as a row-normalized confusion matrix. The compound-level test set was reserved for Module 3 internal testing. For Module 3, performance was evaluated at the compound-level. Internal testing used the test set from the six original compound/control labels; cell-level class probabilities were aggregated within each label to generate six compound assignments, all of which matched the expected class (6/6). External testing used the chemically distinct compounds; cell-level probabilities were aggregated within each compound, and five of seven compounds matched the expected pathway or control class (5/7). For RT-qPCR and other quantitative assays, n denotes the number of independent biological replicates, and qPCR measurements were performed in technical triplicates with GAPDH normalization. For qPCR, two-group comparisons used Welch’s two-sample t-test on log10-transformed fold-change values (*\*P* < 0.05, *\*\*P* < 0.01, *\*\*\*P* < 0.001). Machine-learning accuracy estimates are reported as mean ± SD where repeated runs or resampling estimates were available; uncertainty was estimated by stratified bootstrap resampling within the relevant evaluation set.

## Code availability

The custom code used to train and evaluate the classifier and to reproduce the analyses in this study is available on GitHub at [https://github.com/Pulm-Engine/sigma.git].

## Data availability

Raw imaging flow cytometry images are available from the corresponding author on reasonable request.

## Acknowledgements

The authors thank the UAB Comprehensive Flow Cytometry Core for expert technical assistance and support. This work was supported by National Institutes of Health/National Heart, Lung, and Blood Institute R01HL167010 to Pulin Che and University of Alabama at Birmingham IMPACT Fund.

## Author contributions

P.C., B.G. conceived and supervised the study and acquired funding. X.Z. performed the experiments; M.F. developed the machine-learning framework and performed the computational and statistical analysis. P.C., B.G., X.Z., M.F. prepared the figures and wrote the original draft. All authors reviewed and approved the final manuscript.

